# Frontal and occipito-temporal Theta activity as marker of error monitoring in Human-Avatar joint performance

**DOI:** 10.1101/402149

**Authors:** Q. Moreau, M. Candidi, V. Era, G. Tieri, S.M. Aglioti

## Abstract

Discrepancies between sensory predictions and action outcome are at the base of error coding. However, these phenomena have mainly been studied focusing on individual performance. Here, we explored prediction errors during a human-avatar motor interaction and focused on both the classical frontal error-related brain responses and the activity of the action observation network. Our motor interaction paradigm required healthy individuals to synchronize their reach-to-grasp movements with those of a virtual partner in conditions that did (Interactive) or did not require (Cued) movement prediction and adaptation to the partner’s actions. Crucially, in 30% of the trials the virtual partner suddenly and unpredictably changed its movement trajectory thereby violating the human participant’s expectation. These changes elicited error-related neuromarkers (ERN/Pe - Theta/Alpha modulations) over fronto-central electrodes mainly during the Interactive condition. Source localization and connectivity analyses showed that the frontal Theta activity induced by violations of the expected interactive movements was in phase with occipito-temporal Theta activity. These results expand current knowledge about the neural correlates of on-line motor interactions linking the frontal error-monitoring system to visual, body motion-related, responses.

## Introduction

Interpersonal motor coordination requires dynamic and efficient encoding of others’ actions and spatio-temporal synchronization between individuals (Sebanz et al., 2006; Moreau et al., 2016), thus involving several mechanisms ranging from action perception to goal prediction (Pezzulo, 2013; Panasiti et al., 2017) and motor adaptation (Era et al., 2018; 2019). When interacting without having physical contact with a partner, coordination with the partner’s on-going behaviour is supported by the integration and the prediction of visual and sensorimotor information about one’s own and others’ actions.

However, at times, one’s motor predictions happen to be wrong. The success of adaptive social behaviours relies on the ability to detect prediction errors regarding the movements of the partner and readjust one’s own movement accordingly.

At the neural level, individuals’ ability to predict the fate of observed actions (Aglioti et al., 2008; Abreu et al., 2017) is thought to rely on the activity of the Action Observation Network (AON, Rizzolatti & Craighero, 2004; van Overwalle & Baetens, 2009; Molenberghs et al., 2012; Hardwick et al., 2018) comprising occipito-temporal regions responsible for visual processing of body images (Extrastriate Body Area, EBA; lateral occipito-temporal cortex, LOTC) and biological motion (Superior Temporal Sulcus, STS, Puce & Perrett, 2003; Giese & Poggio, 2003) as well as parietal (anterior Intra Parietal Sulcus, aIPS) and premotor (ventral and dorsal PreMotor, vPM, dPM) areas where the transformation of visual information into motor coordinates is thought to be computed (Keysers & Gazzola, 2014).

The current study focuses on the detection of and adaptation to unforeseen actions from a partner (i.e. prediction errors). Neural correlates of error performance have been previously investigated thoroughly using experimental paradigms such as the Flanker (Hermann et al., 2004) and Simon task (Masaki et al., 2007; Cohen, 2011). EEG studies established that detecting and evaluating our own errors generate two ERPs - the Error-Related Negativity (ERN; Falkenstein et al., 1991; Gehring et al., 1993) and the Positivity error (Pe; Falkenstein et al., 2000) - recorded over fronto-central and parietal electrodes (i.e. FCz and Pz), respectively. The ERN and the Pe are thought to reflect distinctive processes of the monitoring system. It has been proposed that the ERN, which originates from the anterior cingulate cortex (ACC; Carter et al., 1998; van Veen et al., 2001), underlies the detection of “high-level errors” (i.e. failure to meet a goal; Krigolson & Holroyd, 2006), conflict monitoring (Botvinick et al., 2001; Yeung et al., 2004) or action outcome predictions (Quilodran et al., 2008; Alexander & Brown, 2011) while the later and more posterior (centro-parietal) Pe is thought to reflect the detection of “low-level motor errors” (i.e. differences between real and anticipated motor commands, Krigolson & Holroyd, 2007) and is sensitive to error awareness (Endrass et al., 2005; Overbeek et al., 2005). In the time-frequency domain, EEG studies described Theta (4-7Hz) and Alpha (8-13Hz) synchronizations over fronto-central electrodes as markers of the performance monitoring system activity (Luu et al., 2004; Trujillo & Allen, 2007; Cavanagh et al., 2009; Cohen, 2011). A recent study showed that inducing Theta over FCz by means of transcranial alternating current stimulation (tACS) resulted in modulation in behavioural adjustment after error execution (Fusco et al., 2018). More recently, Theta and Alpha synchronizations have also been detected during the observation of motor errors performed by an embodied avatar seen from a first-person perspective (Pavone et al., 2016; Spinelli et al., 2017; Pezzetta et al., 2018).

Crucially, observing someone else performing an error induces similar time and time-frequency EEG responses to those generated when performing an error in first person (van Schie et al., 2004; Miltner et al., 2004; Koelewijn et al., 2008; de Brujin et al., 2007). Thus, the frontal error monitoring system (including the ACC) is considered a generic error processing system which may code in similar ways errors performed by an individual and those observed in another. However, other regions may contribute to error encoding such as inferior parietal areas and insular cortices (Malfait et al., 2010; Orr & Hester, 2012; Shane et al., 2008), as well as occipito-temporal nodes of the AON (Abreu et al., 2012).

Studying error perception during an interactive scenario is central to the broader understanding of the neural correlates of motor prediction, motor control and accurate behavioural responses to unexpected movements during a joint performance. Such studies represent a challenge at a technical level (Moreau & Candidi, 2016), as the simultaneous recording of more than one subject increases the complexity of the experimental design (e.g. increases the number of unexcepted actions within and across participants). Moreover, studying errors in interactive contexts raises relevant theoretical considerations, as the definition of the error (and the related neural responses to it) needs to be framed according to the integration of one’s own and other’s actions and to the outcome at the couple level (i.e. success or failure of the common goal). The latter notion bears on the problem of not knowing whether, and how, the error monitoring system dedicates resources to monitor the action that one needs to perform, the action of the partner and the outcome of the interaction. Furthermore, it is not known whether neural structures beyond the classical AON might be sensitive to others’ errors during an interaction similarly to what happens during observation of others’ actions in non-interactive tasks (Abreu et al., 2012).

To deal with this complexity, we studied modulations of frontal error-related neuromarkers in time and time-frequency domains during an interactive task in which a Virtual Avatar changes its movement trajectory, thus creating a mismatch between what the partner predicts and what he actually sees. Importantly, we also tested whether the lateral occipito-temporal cortex might respond to interactive errors based on the notion that this region may forward visual information to fronto-parietal areas and also may receive top-down information about forthcoming perceptual predictions from frontal nodes of the AON (Keysers & Gazzola, 2015; Zanon et al., 2018). According to predictive accounts of perception (Kilner et al., 2007) top-down (motor) predictions would filter sensory inflow and generate prediction errors in lower level, visual areas (e.g. lateral occipito-temporal cortex) thus implying that frontal error systems would work in phase with occipito-temporal visual areas.

More specifically, we explored in healthy participants whether electrocortical markers of erroneous prediction about some virtual avatar movements specifically code for information needed to accurately perform one’s own action (see Figure 1; Sacheli et al., 2015a; 2015b; Candidi et al., 2017).

**Figure 1.**
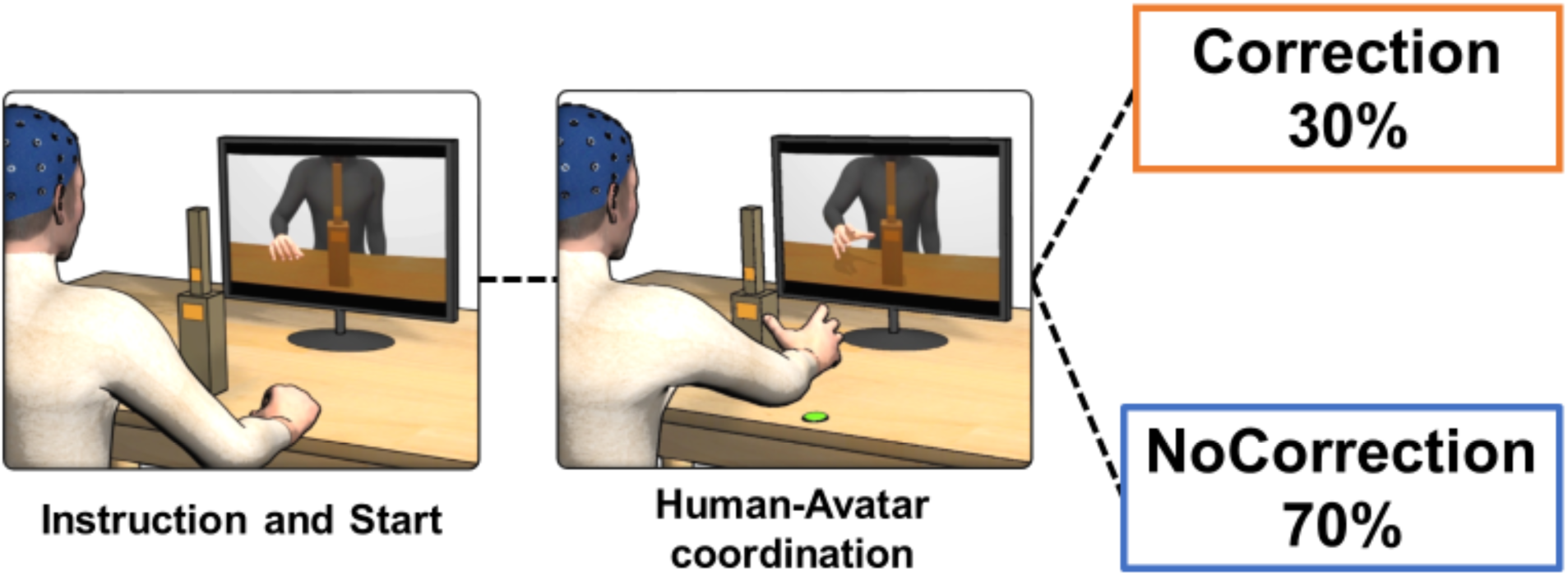
Schematic representation of the experimental set-up.

Participants were asked to reach and grasp a bottle in front of them (with an upper and a lower grasping site) and to synchronize their grasping timing with a virtual partner in two separate conditions, namely: 1) a Cued condition, requiring participants to adapt only the timing of their movements in order to synchronize their touch time with the virtual partner (participants knew in advance where they have to grasp) and, 2) an Interactive condition, requiring participants to adapt in time and space as they needed to coordinate their action according to the avatar’s movement and the instruction received, i.e. imitate or complement the movement of the partner. Crucially, in 30% of the trials, the Virtual Avatar performed an unexpected trajectory change and grasped the other site of the bottle-shape object. This change in the avatar’s movement generated in the human participants the need to adjust their trajectory in reaction to events that diverged from what they had expected (i.e. prediction error). Furthermore, by asking participants to perform imitative or complementary (with respect to those of the virtual partner) reach-to-grasp actions, we explored whether a full match between observed and executed movements would induce any difference in the neural activity associated to the interaction. To this aim, in addition to analysing the classic fronto-central EEG markers, we performed whole-brain analysis to investigate if prediction errors in an interactive motor task would modulate the activity of regions outside the error monitoring system and supporting the visual processing of observed actions.

## Material and Methods

### Participants

22 individuals (13 females, mean age: 26.35 [19-31]; S.D. = 3.54) took part in the experiment. All participants were right-handed with normal or corrected-to-normal vision. Participants were naive as to the aim of the experiment at the outset and were informed of the purpose of the study only after all the experimental procedures were completed. All participants provided written informed consent and were reimbursed 7 €/h. The experimental procedures were approved by the Ethics Committee of the IRCCS Santa Lucia Fondation (Rome, Italy) and the study was performed in accordance with the 2013 Declaration of Helsinki. One participant was detected as an outlier (see below) and therefore removed from all EEG analysis.

### Experimental stimuli and set-up

Participants were comfortably seated in front of a rectangular table of 120 × 75 cm and viewed a 1.024 × 768 resolution LCD monitor placed on the centre of the table at ∼60 cm from their eyes. Participants were asked to reach and grasp a bottle-shaped object (37 cm total height) constituted by two superimposed rectangles with different diameters (small, 2.7 cm; large, 6.5 cm) placed next to the centre of the working surface. To record participants’ grasping time of the bottle, two pairs of touch-sensitive markers (one pair per rectangle) were placed at 15 cm and 22 cm along the vertical height of the object (see Figure 1). Before each trial, participants positioned their right hand on a starting button placed at 34 cm from the bottle-shaped object with their index finger and thumb. The tasks (see below) asked participants to grasp the bottle in synchrony with the avatar appearing on the screen in front of them. Note that the avatar’s index-thumb contact times were measured trial-by-trial by a photodiode placed on the screen that sent a signal recorded by E-Prime 2 professional (version 2.0.10.242, Psychology Software Tools Inc., Pittsburgh, PA) by means of a TriggerStation (BrainTrends ltd., Italy). The photodiode was triggered by a white dot displayed on the screen (not visible to the participants) during the clip frame corresponding to the instant when the avatar grasped its virtual object.

### Creation of the virtual interaction partner

The kinematic features of the virtual partner were based on the movements of human participants performing different grasping movements during a human-human joint-grasping task, identical to the procedures described in Candidi et al. (2017) (see Tieri et al., 2015; Fusaro et al., 2019 for technical details of the Motion Capture recording). The final processed trajectories were realized and applied to a Caucasian male character by using commercial software MotionBuilder 2017 and 3DS Max 2017 (Autodesk). Since we wanted the participants to ignore facial expressions of the virtual partner, the final video stimuli contained only the upper body down from the shoulders, without the neck and head. The complete sample of clips comprised 10 different grasping movements. Half of the movements ended when the hand grasped the bottom part (that is, required power grips Figure 2, Panel A), whereas the other half of the movements ended when the hand grasped the top part of the bottle-shaped object (that is, required precision grips, Figure 2, Panel B). In 30% of the trials the grasps included an online correction, in which the avatar performed a movement correction by switching from a precision to a power grip (or vice versa) during the reaching phase. The correction-videos were created in 3DS Max by merging the initial key frames of a clip (e.g. a power grasp clip) with the last key frames of a different clip (e.g. precision grasp clip) (Figure 2, Panel C-D).

**Figure 2.**
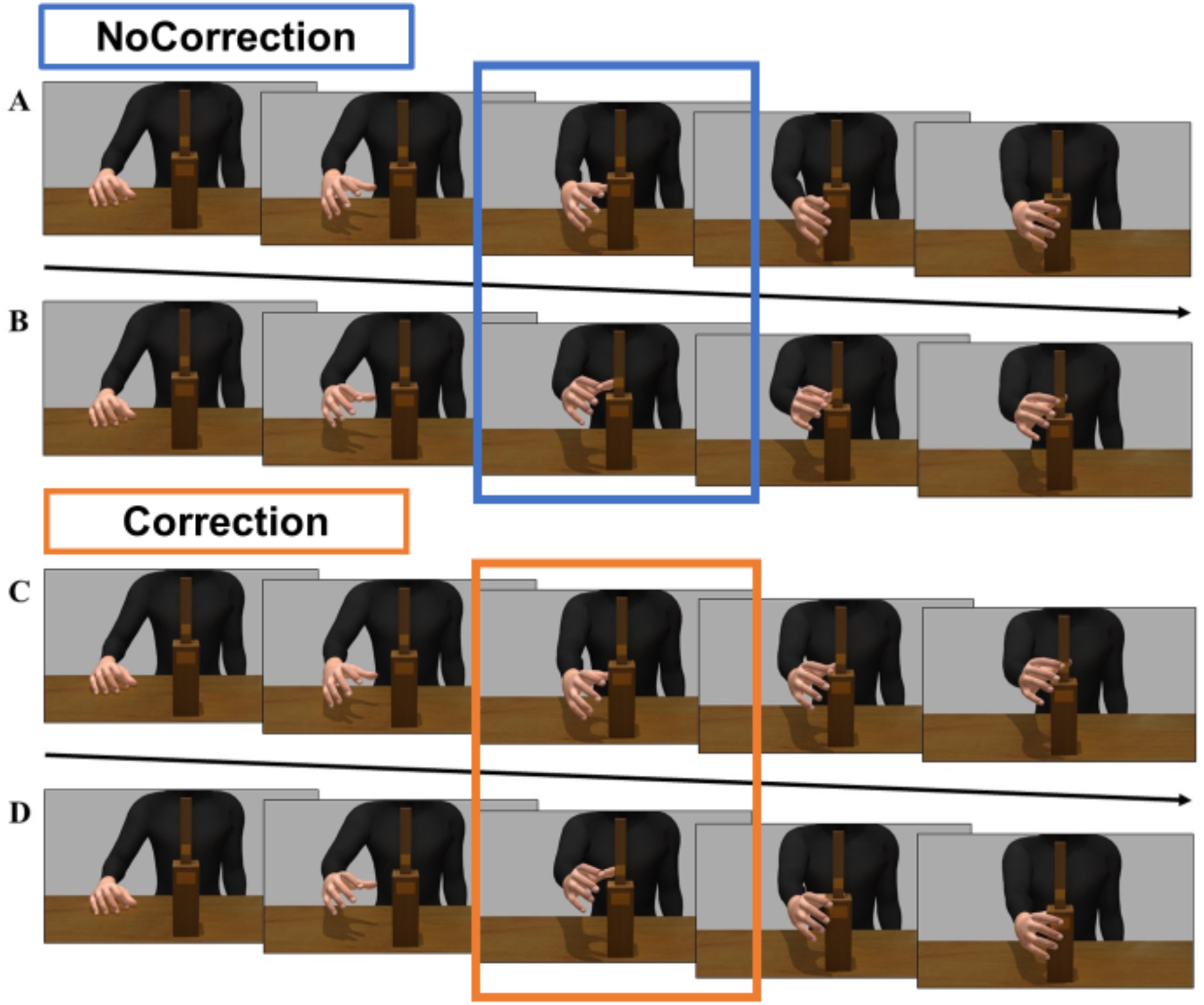
Examples of the sequence of frames for each type of virtual partner’s movements: A) Power Grasp; B) Precision Grasp; C) Correction Power to Precision and D) Correction Precision to Power Grasp; The middle frame of each sequence represents the 0 time point for EEG marker in which the Avatar corrects (orange) or does not correct (blue) its behaviour.

### Experimental Task

We used an ecological and controlled human-avatar interactive task (Sacheli et al., 2015a; 2015b; 2018; Candidi et al., 2017; Era et al., 2018a; Gandolfo et al., 2019), which has been shown to recruit the same behavioural processes called into play during human-human interaction, namely mutual adjustment and automatic imitation (Sacheli et al., 2012; 2013; Candidi et al., 2015; Curioni et al., 2017; Era et al., 2018b). Importantly, in the present task, one’s own action goal cannot be achieved without considering the virtual partner’s online movements and adapting to them. Participants were required to reach and grasp the bottle-shaped object placed in front of them with their right hand, as synchronously as possible with the action of the avatar (shown on the screen in front of them) in respect to its bottle-shaped object. Given the dimensions of the bottle-shaped object, grasping the lower part implied a whole-hand grasping (a power grip), whereas grasping the upper part implied a thumb-index finger precision grip (Movement Type Factor).

Participants performed the task in two different conditions (Condition Factor): (1) the “Cued Condition”, where subjects received either a high pitch sound (indicating that they had to grasp the bottle in the upper part) or a low pitch sound (indicating that they had to grasp the bottle in the lower part), and (2) the “Interactive Condition”, where subjects received either a sound indicating that they had to perform an imitative action (i.e. participant and virtual partner both grasping the upper or lower part of their bottle) or a sound indicating they had to perform a complementary action (i.e. if the virtual partner was grasping the lower part of its bottle, participant had to grasp the upper part of his/her bottle, and vice versa). The timeline of a trial was as follows: participants received the auditory instruction about what kind of action they should perform, followed by the presentation of a fixation cross (300ms), preceding the appearance of the Avatar on the screen; the Avatar started its movement between 200 and 600ms after a “go signal” was given and performed its action (trial duration ∼2000 ms). The inter-trial interval depended on the time participants took to go back from the bottle to the starting position. The experimenter manually triggered the next trial as soon as participants went back to the starting position.

In the Cued Condition, participants had to predict and adapt to the avatar’s movement in time (i.e. when the virtual partner is going to grasp the bottle) but not in space, since they knew in advance where they had to grasp the bottle-shaped object, whereas, in the Interactive Condition, participants had to predict and adapt in time *and* space (i.e. when and where the virtual partner is going to grasp the bottle). It was emphasized that in all the conditions participants had to touch the bottle as synchronously as possible with the virtual partner.

The clips’ frame during which the avatar started correcting its behaviour (e.g. by switching from a power to a precision grasp or vice versa, Correction Factor) was used as the 0-time-point for EEG markers. In the trials where the virtual partner did not correct its behaviour, the time 0 corresponds to the same frame where the switching would have happened in the change clip (see above, Figure 2). Participants performed four 100-trial blocks (2 blocks of the Cued Condition, 2 for the Interactive Condition, presented in a counterbalanced order between participants). In 30% of the trials, the virtual partner performed a correction. Thus, each participant performed in 140 trials for Cued-NoCorrection, 140 trials for Interactive-NoCorrection, 60 trials for Cued-Correction and 60 trials for Interactive-Correction. The interaction type (Complementary/Imitative) and the movement type (Precision/Power) factors were randomized trial-by-trial and pooled together. While an unequal number of Correction and No-Correction trials is usually used to emphasize the salience of one condition with respect to another (Pavone et al., 2016) we demonstrated that randomly selecting an equal number of trials for the two distributions does not change the results (Pezzetta et al., 2018). Stimuli presentation and randomization were controlled by E-Prime2 Professional software (Psychology Software Tools Inc.).

### Behavioural data

We considered the Grasping Synchrony as the main behavioural measure, i.e. the absolute value of the time delay between subjects’ index-thumb contact-times on their bottle and the avatar’s bottle touch time (Sacheli et al., 2015a). This showed the success of human-avatar coordination. Analysis regarding other behavioural and kinematics measures are reported as Supplementary Materials.

### EEG preprocessing

EEG signals were recorded and amplified using a Neuroscan SynAmps RT amplifiers system (Compumedics Limited, Melbourne, Australia). These signals were acquired from 60 tin scalp electrodes embedded in a fabric cap (Electro-Cap International, Eaton, OH), arranged according to the 10-10 system. The EEG was recorded from the following channels: Fp1, Fpz, Fp2, AF3, AF4, F7, F5, F3, F1, Fz, F2, F4, F6, F8, FC5, FC3, FC1, FCz, FC2, FC4, FC6, T7, C5, C3, C1, Cz, C2, C4, C6, T8, TP7, CP5, CP3, CP1, CPz, CP2, CP4, CP6, TP8, P7, P5, P3, P1, Pz, P2, P4, P6, P8, PO7, PO3, AF7, POz, AF8, PO4, PO8, O1, Oz, O2, FT7 and FT8. Horizontal electro-oculogram (HEOG) was recorded bipolarly from electrodes placed on the outer canthi of each eye and signals from the left earlobe were also recorded. All electrodes were physically referenced to an electrode placed on the right earlobe and were algebraically re-referenced off-line to the average of both earlobe electrodes. Impedance was kept below 5 KΩ for all electrodes for the whole duration of the experiment, amplifier hardware band-pass filter was 0.01 to 200 Hz and sampling rate was 1000 Hz. To remove the blinks and eyes saccades, EEG and horizontal electro-oculogram were processed in two separate steps. Data were then downsampled at 500 Hz before a blind source separation method was applied on continuous raw signal, using Independent Component Analysis (ICA) (Jung et al., 2000) implemented in the Matlab toolbox EEGLab_(version 14_1_1_b running on Matlab 2010a) (Delorme & Makeig, 2004) to remove any components related to eye movements from the EEG. 59 components were generated and artifactual components (blinks and saccades) were removed based on the topography and the explained variance (components are ordered by the amount of variance they represent), data were then visually inspected after the removal of the components, to check if both blinks and saccades were no longer present (2.83 components per participants were rejected on average). The signal was then segmented into epochs of 2000 ms (−1000 ms to +1000 ms around the Avatar’s Correction/NoCorrection frame) and visually inspected to control for residual eyes movements as well as muscular artifacts. Bad epochs were identified and removed from further analysis; the artifact rejection procedure led to 9.2% of the trials being rejected. The average number of trials per participants was 127.9 trials for Interactive-NoCorrection, 128.4 trials for Cued-NoCorrection, 56.5 trials for Cued-Correction and 55.5 trials for Interactive-Correction. For all EEG variables presented below, participants with a mean 2.5 SDs above or below the group mean were excluded from the analyses. According to this criterion, one participant was detected as an outlier for the Theta ERD/ERS and therefore removed from all EEG analyses, resulting in 21 kept participants for all analyses.

### EEG Analysis

#### ERPs

Time domain analyses were performed by using the FieldTrip (version 2015-01-13) routines (Donders Institute, Nijmegen; Oostenveld et al., 2010) in Matlab2010a (The MathWorks, Inc.). The EEG time series were band-pass filtered (2 to 30 Hz) to reduce the contribution of slow potentials that masked some of the frontal components relevant to our study (Pavone et al., 2016). It is held that high-pass filters > 1 Hz may generate artefactual effects in ERPs (Tanner et al., 2015). However, we checked that grand average waveforms with and without filters maintain the same morphology (Acunzo et al., 2012), and did not introduce distortions that may bias the estimated parameters (Widmann et al., 2015). Each epoch was baseline corrected from 200ms to 0ms before the Avatar’s correction (or absence of correction). Two main components, already described in the Error-related ERP literature (i.e., ERN over FCz and Pe over Pz) were individuated. Visual inspection of the results showed the generation of an ERN component only for the Interactive-Correction and Cued-Correction trials (see Figure 3). It also appears that the ERN component peaked at different times for Interactive and Cued conditions (i.e 194 ms for Interactive-Correction and 228 ms for Cued-Correction). Therefore, we extracted the ERP mean amplitude over a time window of 100 ms around the ERNs respective latency-peaks (Spinelli et al., 2018). The Pe component was also identified for Interactive-Correction trials, peaking at 402 ms after Avatar’s correction and peaking at 536 ms for Cued-Correction trials; therefore, we extracted the mean amplitude from a time window of 100ms around these two peaks. As no clear peaks has been detected for Interactive-NoCorrection and Cued-NoCorrection conditions, we extracted the mean amplitudes using the same latencies as the ones used for their respective Correction trials.

**Figure 3.**
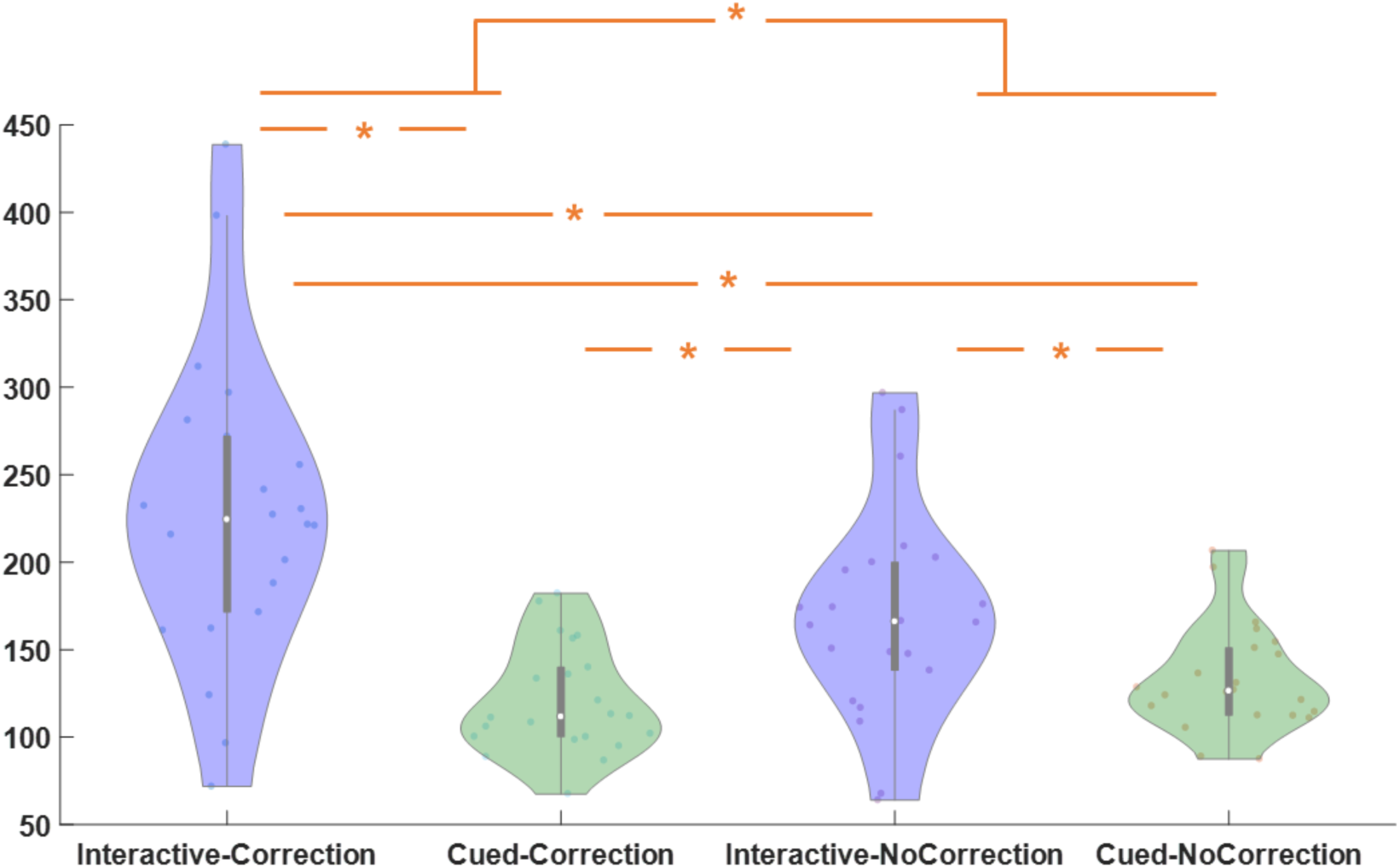
Grasping Synchrony. The ANOVA showed a significant interaction between Correction and Condition (F(1,20)= 49.53, p < 0.001, ηp² = 0.71). The post-hoc test indicated that the synchrony performance was worse for Interactive-Correction compared to the other conditions (all ps < 0.001) and for Interactive-NoCorrection compared to the Cued-Correction and Cued-NoCorrrection conditions (all ps < 0.001). Asterisks indicate significant (p < 0.05) differences.

### ERD/ERS

Time-frequency analyses were performed by using the Fieldtrip (version 2015-01-13) routines (Donders Institute, Nijmegen; Oostenveld et al., 2010) in Matlab2010a (The MathWorks, Inc.). The EEG time series were obtained by segmenting the signal into epochs of 2000 ms length and band-pass filtered (0.1 to 100 Hz). Each epoch was transformed in the frequency using Hanning-tapered window with 4 cycles and a 50 ms time resolution (using the ‘ft_freqanalysis’ function with ‘mtmconvol’ method as implemented in FieldTrip). Estimated frequency band results were displayed as event-related desynchronization/synchronization (ERD/ERS) with respect to a baseline of between −500 and 0 ms (cfg.baselinetype = ‘relchange’) before the Avatar’s change. Positive and negative ERD/ERS values index synchronization and desynchronization with respect to a given reference interval (Pfurtscheller & Lopes da Silva, 1999). For each experimental condition, ERD/ERS were computed from zero (Avatar’s change) to 500 ms. In line with previous literature on frequency modulation during error processing (Cohen, 2011; Cavanagh et al., 2009) we extracted ERD/ERS for the Theta band (4-7 Hz) and the Alpha band (8-13 Hz) and analysed the modulation of power over FCz, see Supplementary Materials for analysis on the Beta band.

### Source Analysis

Beamformer analyses were performed to estimate cortical sources of the effects found at the sensor level and were accomplished using the Dynamic Imaging of Coherent Sources (DICS) approach, as implemented in Fieldtrip to account for frequency specific effects. The cross spectral density matrix was calculated at the frequency of interest (i.e. 5 Hz for the Theta band; 10Hz for the Alpha band). The head model used to project the estimated source was based on a standard MRI template (Holmes et al., 1998; Oostenveld et al., 2003) and the electrodes position used was based on the international standard 10-10 system. Sources activity post-trigger (0-500ms) was contrasted to source activity pre-trigger (−500 to 0 ms). The change in oscillatory power was averaged out cross participants (see Figure 6) and plotted using the Connectome Workbench provided by the Human Connectome Project (Van Essen, 2012; Seymour et al., 2017; see also the shared Matlab/Fieldtrip code “get_source_power.m” at https://github.com/neurofractal/sensory_PAC). Two Region of Interest (ROIs) were identified: a fronto-central and right occipito-temporal one (i.e. right LOTC and Fronto-central ROIs) and average power for these two ROIs was extracted for analysis (see Figure 6).

### Connectivity Analysis

Based on the visual inspection and statistical results of the source analysis, we focused on the two separate source estimates (namely right LOTC and Fronto-central ROI). Using the maximum power coordinates of these two sources (estimated on the subjects’ grand average), we performed Linear Constrained Minimum Variance (LCMV) source analysis in the time domain to extract time series at the two locations of interest, creating two “virtual channels” (i.e. “Fronto-central” and “right LOTC”, see Figure 7). For a description of the entire procedure, see the “MEG virtual channels and seed-based connectivity” tutorial on the Fieldtrip webpage (http://www.fieldtriptoolbox.org/tutorial/chieti/virtualchannel). Once extracted, the virtual channels were treated as normal EEG data and averaged in the time domain (see Figure 7) (Lappe et al., 2013; Baumgarten et al., 2015). Then, we used the complex time-frequency estimates of the two virtual channels to compute the Phase-Locking Value (PLV) between the two regions of interest (See Figure 7B) where the PLV is computed as a value between 0 and 1 that quantifies the phase consistency across multiple trials. The PLV is the absolute value of the mean phase difference between the two signals, expressed as a complex unit-length vector (as described by Lachaux et al., 1999). PLV data were extracted from 300-600ms after the Avatar’s correction in frequencies between 4-11 Hz, based on visual inspection of the data.

### Data handling and Statistics

Our main hypothesis concerns ERPs (ERN, Pe) and time-frequency (Theta, Alpha ERD/ERS) modulations in conditions during which participants need to: i) predict the action of their partner and proactively adapt to it (Interactive/Cued factor); ii) predict and adapt to an error performed by their partner (Correction/NoCorrection factor). Therefore, the analyses presented in the main text focus on these two factors. Moreover, collapsing Interaction Type (Complementary/Imitative) and Movement type (Power/Precision grasping) factors allowed us to have a higher number of trials for each condition. See Supplementary Material for analysis using all the factors.

Grasping Synchrony, ERN and Pe components, Time-frequency indexes (Theta, Alpha ERD/ERS) and the PLV results were analysed through separated 2 x 2, within-subject, repeated measures ANOVA, with Correction (Correction/NoCorrection) and Condition (Interactive/Cued) as within-subject factors.

Source power indexes for the 2 ROIs were analysed through a 2 x 2 x 2 repeated measures ANOVAs with ROIs (Fronto-central/right-LOTC) Correction (Correction/NoCorrection) and Condition (Interactive/Cued) as within-subject factors.

Frequentist statistical analyses (Shapiro-Wilk test for normality, General Linear Model (GLM) and Greenhouse-Geisser correction for non-sphericity when appropriate (Keselman & Rogan, 1980)) were performed with Statsoft Statistica 8 software. Post-hoc correction for multiple comparisons was made using the Bonferroni test. Violin plots have been computed using the shared Matlab function ‘violin..m’ (https://github.com/bastibe/Violinplot-Matlab/blob/master/Violin.m).

## Results

### Behavioural

#### Grasping Synchrony

The 2 Correction (Correction/NoCorrection) x 2 Condition (Interactive/Cued) ANOVA showed that the factors Correction and Condition reached statistical significance with worse synchronicity performance for the Correction vs. the NoCorrection trials (F(1,20) = 15.53, *p* < 0.001, ηp² = 0.43) as well as for Interactive condition vs. the Cued one (F(1,20) = 29.75, *p* < 0.001, ηp² = 0.60). The interaction between Correction and Condition was also significant (F(1,20)= 49.53, *p* < 0.001, ηp² = 0.71). The post-hoc tests indicated that the synchrony performance was worse for Interactive-Correction compared to the other conditions (all *ps* < 0.001), and for Interactive-NoCorrection compared to the Cued-Correction and Cued-NoCorrrection conditions (all *ps* < 0.001) (see Figure 3).

### ERPs

#### ERN over FCz

The 2 Correction (Correction/NoCorrection) x 2 Condition (Interactive/Cued) ANOVA showed that the factors Correction and Condition reached statistical significance as main effects, with larger ERN amplitude for Correction compared to NoCorrection trials (F(1,20) = 31.45, *p* < 0.001, ηp² = 0.61) and larger ERN amplitude during the Interactive condition compared to the Cued one (F(1,20) = 4.46, *p* = 0.47, ηp² = 0.18). The interaction between Correction and Condition also reached statistical significance (F(1,20) = 7.82, *p* = 0.011, ηp² = 0.28). Post-hoc test indicated that ERN amplitude observed during Interactive-Correction trials was larger than the one recorded during all the other conditions (all *ps* < 0.015) and that the ERN amplitude observed for Cued-Correction condition was larger than both Interactive-NoCorrection (*p* < 0.001) and Cued-NoCorrection (*p* = 0.014) (see Figure 4).

**Figure 4.**
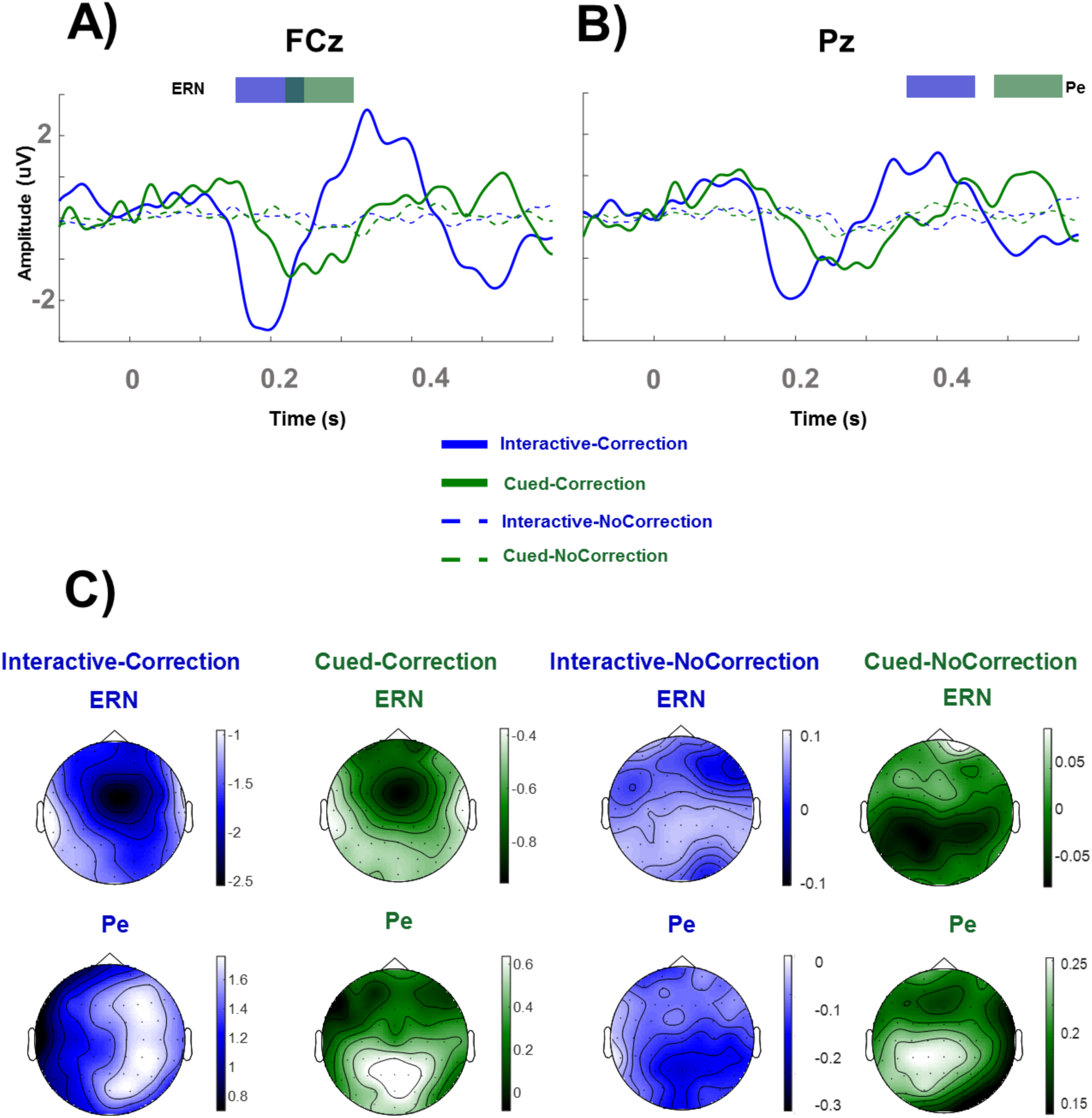
Grand averages of A) the ERN over FCz (144-244 ms for the Interactive conditions, 178-278 ms for the Guided conditions, shown in blue and green thick lines above the Amplitude panel) and B) the Pe over Pz (352-452 ms for the Interactive conditions, 486-586 ms for the Guided conditions, shown in blue and green thick lines above the Amplitude panel) components in the four different experimental conditions, C) Topographies of each components.

#### Pe over Pz

The 2 Correction (Correction/NoCorrection) x 2 Condition (Interactive/Cued) ANOVA showed that the factor Correction reached statistical significance as a main effect (F(1,20) = 25.09, *p* < 0.001, ηp² = 0.55) (see Figure 4).

### Time Frequency

#### Theta (4-7Hz) ERD/ERS Over FCz

The 2 Correction (Correction/NoCorrection) x 2 Condition (Interactive/Cued) ANOVA showed that the factors Correction and Condition reached statistical significance as main effects, with larger Theta synchronization for Correction compared to NoCorrection trials (F(1,20) = 49.61, *p* < 0.001, ηp² = 0.71) and larger Theta synchronization during the Interactive condition compared to the Cued one (F(1,20)= 93.85, *p* < 0.001, ηp² = 0.82). The interaction between Correction and Condition also reached statistical significance (F(1,20) = 20.309, *p* < 0.001, ηp² = 0.50). Post-hoc test indicated the following: 1) Theta ERS during Interactive-Correction trials was larger than the one recorded during all the other conditions (all *ps* < 0.001); 2) Theta ERS in Interactive-NoCorrection condition was larger than Cued-NoCorrection (*p* = 0.001); 3) Theta ERS for Cued-Correction trials was bigger than Cued-NoCorrection trials (*p* = 0.037) (see Figure 5).

**Figure 5.**
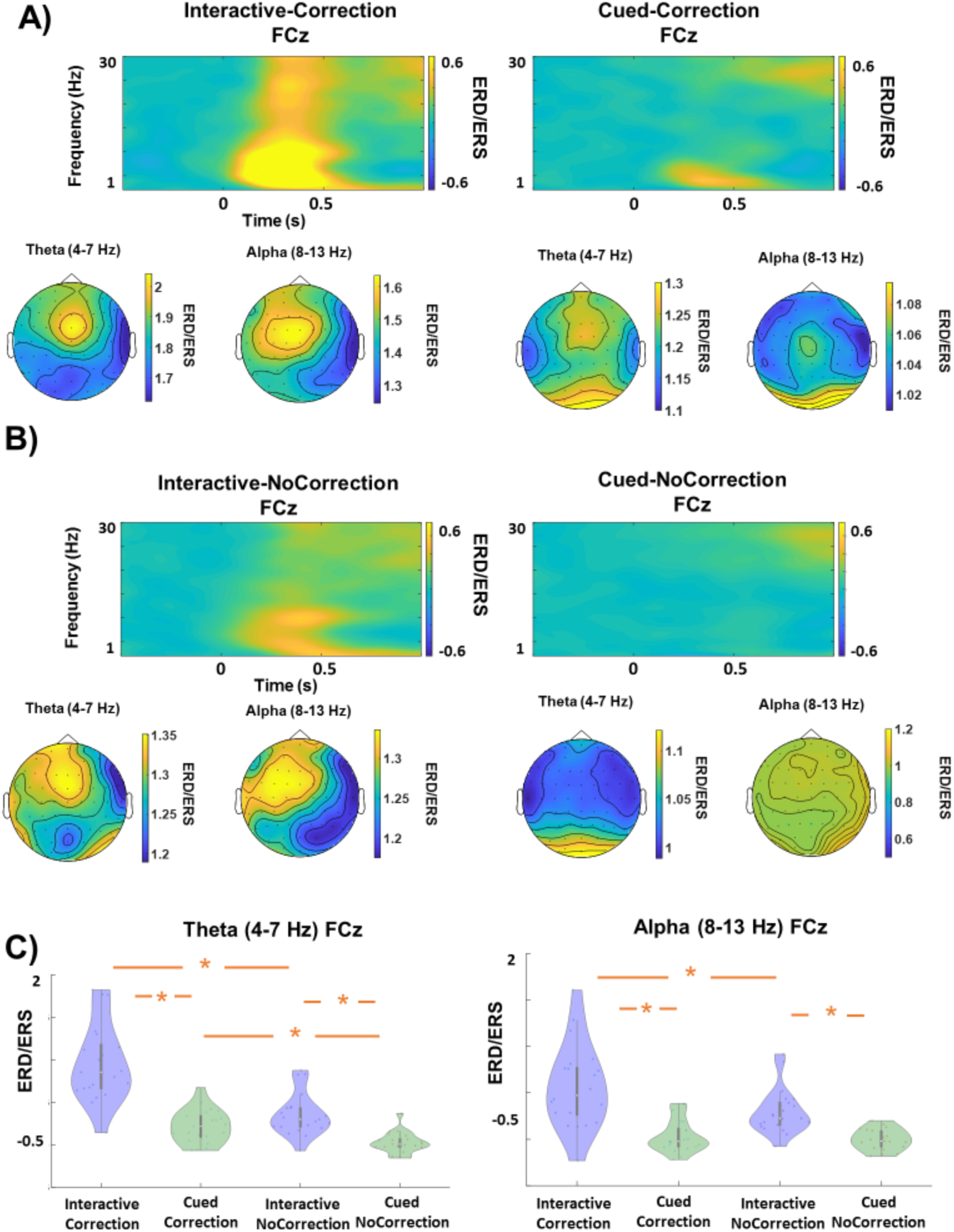
A) ERD/ERS plot (0-30 Hz) over FCz and Theta and Alpha topographical (0-500 ms) views when the Avatar corrected its movement (Correction factor); B) ERD/ERS plot (0-30 Hz) over FCz and Theta and Alpha topographical (0-500 ms) views when the Avatar did not correct its movement (NoCorrection factor); C) The left plot shows the Theta ERS at FCz and the right plot shows the Alpha ERS at FCz (see text for the description of the effects).

Previous literature has highlighted the common origin of the ERN and Theta synchronization (Luu et al., 2004; Trujillo & Allen, 2007). To assess this matter, we ran a correlation between the ERN mean amplitude and the Theta ERS (see Table 1). The analysis shows a significant negative correlation (due to negative amplitude of the ERN) between Theta and ERN over FCz for both Interactive-Correction and Cued-Correction conditions.

**Table 1.**
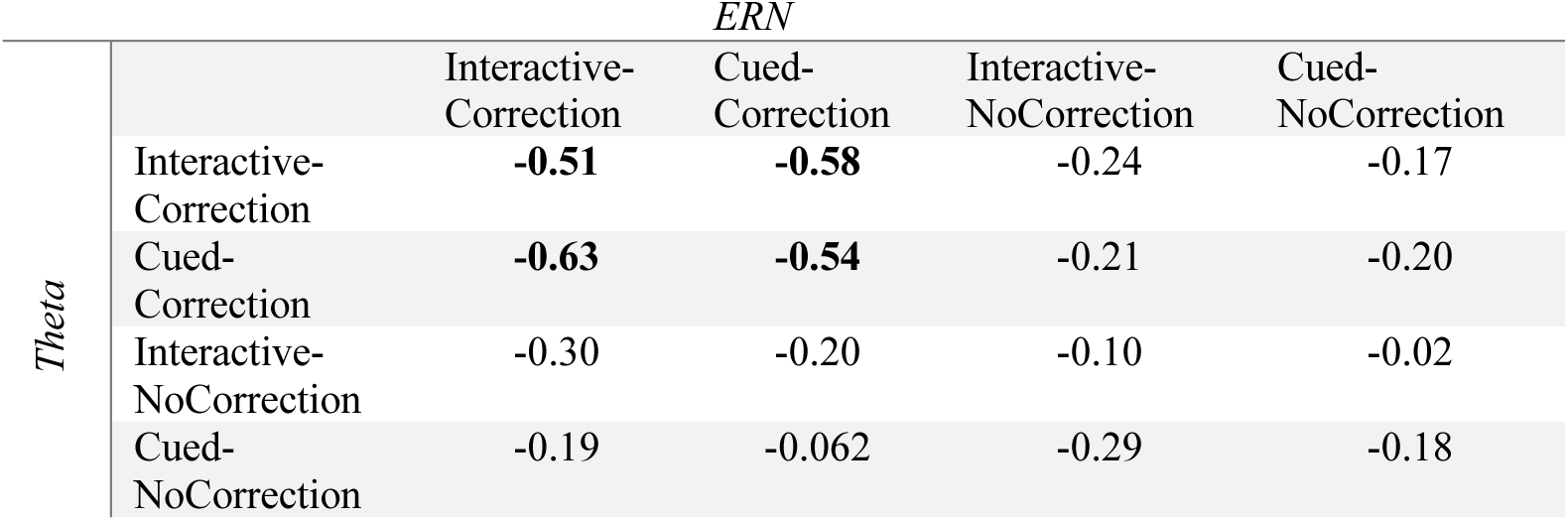
Correlations between Theta and ERN over FCz, bold values shows significant correlation (*p* < 0.05).

#### Alpha (8-13Hz) ERD/ERS Over FCz

The 2 Correction (Correction/NoCorrection) x 2 Condition (Interactive/Cued) ANOVA showed that the factors Correction and Condition reached statistical significance as main effects, with larger Alpha synchronization for Correction compared to NoCorrection trials (F(1,20) = 13.20, *p* = 0.002, ηp² = 0.39) and larger Alpha synchronization during the Interactive condition compared to the Cued one (F(1,20) = 35.71, *p* < 0.001, ηp² = 0.64). The interaction between Correction and Condition also reached significance (F(1,20) = 10.99, *p* = 0.008, ηp² = 0.30). Post-hoc test indicated that the Alpha ERS for Interactive-Correction was bigger than those generated by all the other conditions (all *ps* < 0.001), and that Alpha ERS for Interactive-NoCorrection was bigger than that recorded during Cued-NoCorrection trials (*p* < 0.001) while no difference was shown between Cued-Correction and Cued-NoCorrection (*p* = 1) (see Figure 5).

To sum-up, both Theta and Alpha ERS are influenced by the different conditions. In details, there is stronger Theta and Alpha synchronization for Interactive-Correction trials compared to all other conditions, and stronger Theta and Alpha synchronization for Interactive-NoCorrection compared to Cued-NoCorrection trials. Frontal midline Theta is usually associated with error processing, but it has been shown that Theta power could affect nearby frequencies (i.e. Delta and Alpha; Trujillo & Allen, 2007). To determine if the Alpha activity over FCz reflects a spreading of the Theta activity, which is stronger than the Alpha one in absolute value, we ran a correlation between Theta-ERS and Alpha-ERS (see Table 2). The analysis shows a significant positive correlation between Theta and Alpha power over FCz for each condition (matrix diagonal). Topographical similarities, similar patterns of results and significant correlations suggest a possible spreading of the Theta band over the Alpha-band.

**Table 2.** Correlations between Theta and Alpha ERS over FCz, bold values shows significant correlation (*p* < 0.05).

|  |  | Theta |  |  |  |
| --- | --- | --- | --- | --- | --- |
| Alpha |  | Interactive-Correction | Cued-Correction | Interactive-NoCorrection | Cued-NoCorrection |
|  | Interactive-Correction | 0.61 | 0.59 | 0.36 | 0.28 |
|  | Cued-Correction | 0.09 | 0.68 | -0.04 | 0.40 |
|  | Interactive-NoCorrection | 0.39 | 0.10 | 0.66 | 0.23 |
|  | Cued-NoCorrection | -0.03 | 0.72 | -0.01 | 0.70 |

### Source Analysis

Two Regions of Interest (ROIs) were identified for Theta and Alpha power modulations: a fronto-central and right occipito-temporal one (i.e. right LOTC and Fronto-central ROIs) and average power for these two ROIs was extracted for analysis (see Figure 6).

**Figure 6.**
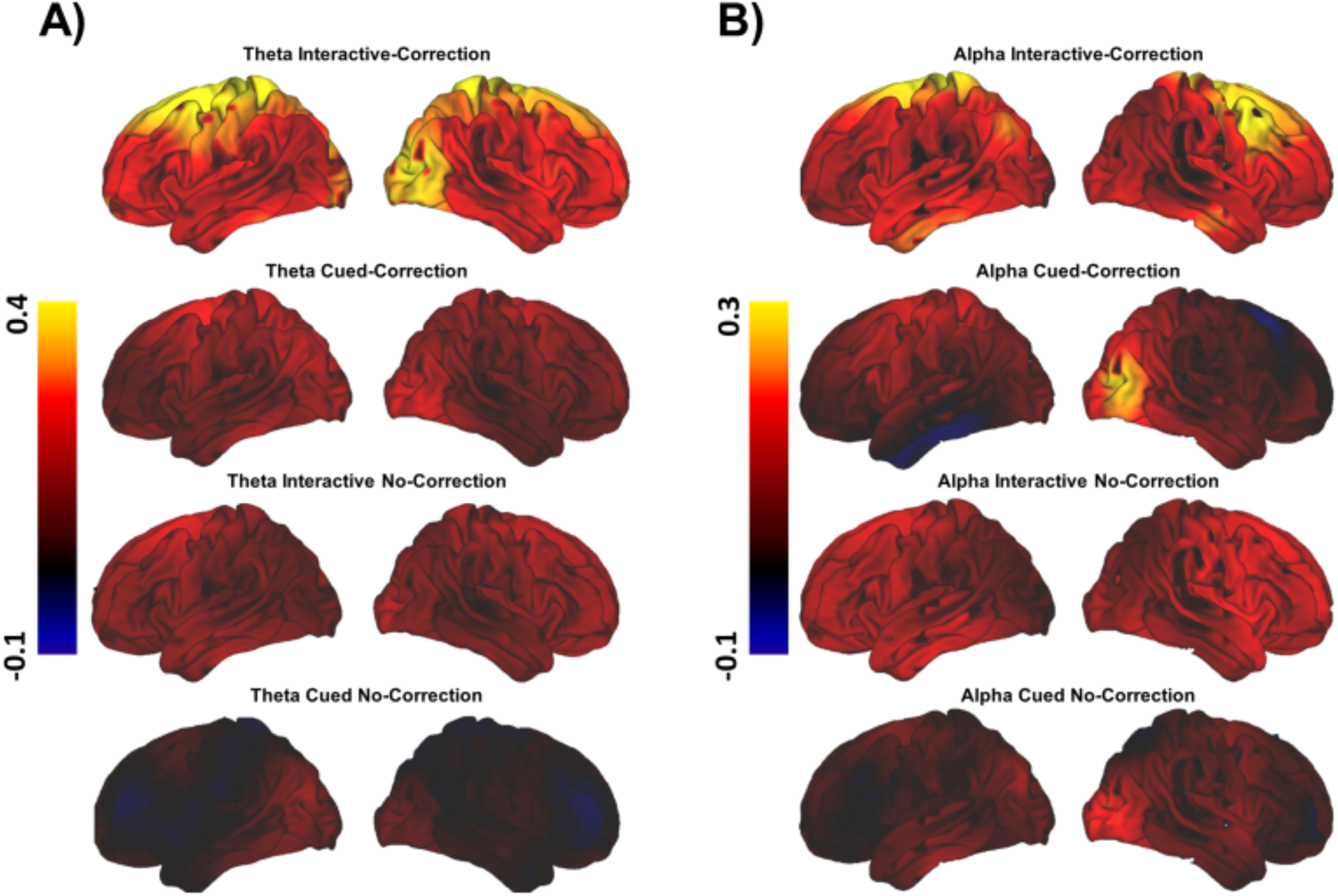
Whole-brain A) Theta and B) Alpha source power.

#### ANOVA on ROIs - Theta Source – 0-500ms post observed error

The 2 ROIs (Fronto-central/right-LOTC) x 2 Condition (Interactive/Cued) x 2 Correction (Correction/NoCorrection) ANOVA showed that both the factor Condition reached statistical significance as a main effect, showing more Theta source power for Interactive compared to Cued trials (F(1,20) = 16.64, *p* < 0.001, ηp² = 0.45), and the factor Correction showing more Theta source power for Correction compared to Non-Correction trials (F(1,20) = 57.38, *p* < 0.001, ηp² = 0.74). Two interactions also reached significance: 1) the interaction between Condition and Correction (F(1,20) = 6.91, *p* = 0.016, ηp² = 0.25) showing that the Interactive-Correction condition showed more Theta power than all the other conditions (*ps <* 0.001), and that the Cued-NoCorrection condition generated less Theta source power than Cued-Correction and Interactive-NoCorrection conditions (*ps <* 0.001); 2) the interaction between ROIs and Condition (F(1,20) = 4.48, *p* = 0.046, ηp² = 0.18) showing that the both ROIs in the Interactive condition showed more Theta activity than both ROIs in the Cued condition (*ps <* 0.002).

#### ANOVA on ROIs - Alpha Source – 0-500ms post observed error

The 2 ROIs (Fronto-central/right-LOTC) x 2 Condition (Interactive /Cued) x 2 Correction (Correction /NoCorrection) ANOVA showed that both the factor Condition reached statistical significance as a main effect, showing more Alpha source power for Interactive compared to Cued trials (F(1,20) = 21.70, *p* < 0.001, ηp² = 0.52), and the factor Correction showing more Alpha source power for Correction compared to Non-Correction trials (F(1,20) = 91.38, *p* < 0.001, ηp² = 0.82). No interaction reached significance (*ps* < 0.08).

In sum, the Fronto-central and the right-LOTC ROIs show similar patterns, namely a significant increase of Theta and Alpha source power during the Interactive-Correction condition compared to Cued-Correction (Figure 6). By using the coordinates of these two ROIs, we subsequently targeted functional connectivity between these two areas.

### Fronto-central-right occipito temporal connectivity - Phase-Locking Value

The 2 Correction (Correction/NoCorrection) x 2 Condition (Interactive/Cued) ANOVA showed that the factors Correction and Condition reached statistical significance as main effects, with larger Phase Locking Value for Correction compared to NoCorrection trials (F(1,20) = 57.93, *p* < 0.001, ηp² = 0.74) and larger PLV during the Interactive condition compared to the Cued one (F(1,20) = 9.53, *p* = 0.005 ηp² = 0.32) (see Figure 7). These results suggest an increase in phase-locking between the Fronto-central ROI and the right LOTC in the 4-11Hz frequency range during the Interactive condition and when the Avatar corrected its movement.

**Figure 7.**
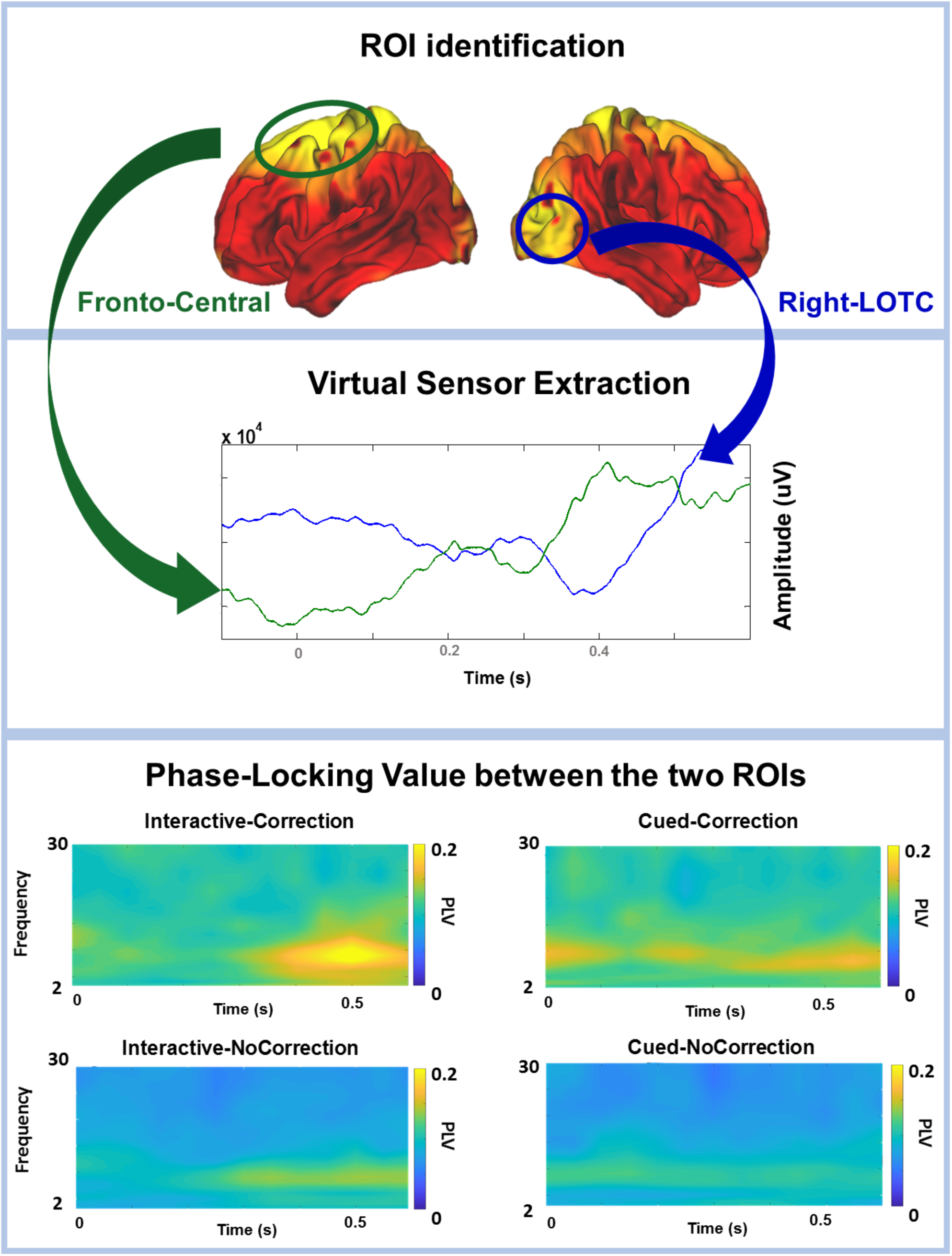
Functional connectivity between the two virtual sensors (Fronto-central and Right-LOTC).

## Discussion

In the present study we recorded EEG in participants who performed a human-avatar joint-grasping task to explore the link between visual action prediction and action monitoring systems in an interactive context where a virtual partner could perform actions that violated the motor prediction of human participants. We obtained four main results: 1) electrocortical indices of error monitoring were higher in conditions requiring the participant to predict in space and time the outcome of their partner’s action (Interactive condition) compared to when only coordination in time was required (Cued condition); 2) modulation of the above-mentioned indices, particularly of theta activity over fronto-central electrodes, was stronger in conditions where the virtual partner changed its initial action (Interactive-Correction condition); 3) the virtual partner’s correction generates an additional increase of occipito-temporal theta activity; 4) there is an increased frontal and occipito-temporal theta band connectivity both when the avatar unpredictively changed its movement (Correction trials) and when the actions of the participants depend on those of the virtual partner (Interactive condition).

### Action and error monitoring during motor interactions

Studies show that similar activity is found when people perform errors (Debener et al., 2005; Gehring et al., 1993) and observe another person making an error (van Schie et al., 2004; Malfait et al., 2010; Cracco et al., 2016; Desmet & Brass, 2015). This suggests that the detection of ones’ own error and of those made by others shares analogous neural mechanisms. Furthermore, individuals’ motor expertise in specific action domains influences behavioural and neurophysiological responses to erroneous action observation (Aglioti et al., 2008; Abreu et al., 2012; Candidi et al., 2012; Panasiti et al., 2016) suggesting that monitoring one’s own actions and the actions of others is an inherently plastic process.

Besides the suggested overlap between the neural systems responding to observed and executed actions and errors (Zubarev et al., 2018) studies have documented a differential contribution of brain areas to the observation of errors performed by others (Shane et al., 2008; Abreu et al., 2012; Somon et al., 2019; Ninomiya, et al., 2018; Somon et al., 2017). In the present study we explored the neural responses associated to coordinating one’s own movements with those of a virtual partner who could perform unexpected changes in its motor behaviour. When in a dyad, both members need to fulfil their own motor sub-goal while aiming at coordinating each other’s actions in order to achieve a common goal.

In all the experimental conditions of the present study, the individual goal is to grasp a target in synchrony with a partner. Crucially, however, in the Interactive condition participants need to realize imitative or complementary interactions which can only be achieved by taking into account both one’s own and the avatar’s behaviour. In this sense, the definition of an error in the Interactive condition pertains to linking self-other action perception and execution. Hence, participants had to monitor visual inputs from the body of the partner to plan their own action. We suggest that the higher theta phase locking between frontal-error-related and occipito-temporal (i.e. visual cortices) areas in the Interactive condition illustrates this phenomenon. We speculate that during interactive scenarios, given the bodily nature of the visual information, visual nodes of the AON may support the activity of the error monitoring system to allow behavioural adaptation.

### Error-related ERPs

Error-Related-Negativity is usually associated with an early detection of an unexpected outcome (compared to an internally-generated prediction) which may be represented in our study by the unpredicted Avatar’s movement. The ERN over FCz reveals specific modulation of error monitoring associated with Interactive-Correction trials. The components were only visually identified in conditions where the Avatar changed its behaviour (Correction factor) (see Figure 4). Interestingly, the Avatar’s changes in the Interactive condition elicited greater ERN mean amplitudes than in the Cued condition, while the Pe was only modulated by the Correction factor. This pattern of results indicates that time-dependent neural responses triggered by error detection are induced by others’ errors, and that the ERN is modulated by the relevance of these errors in one’s own movements during interaction.

Besides the relevance of the others’ error, a recent study found that the ERN is also influenced by the magnitude of an observed error in space with greater amplitude and earlier latency for large errors compared to small ones (Spinelli et al., 2018). An imaging study complemented this error-magnitude pattern of neural responses showing that also activity in occipito-temporal cortex is sensitive to the magnitude of observed reaching deviations (Malfait et al., 2010). Furthermore, in Table 1 we show correlations between ERN amplitude and Theta ERS, only in Corrected conditions (where ERN is present). This is in line with previous research claiming a link between phase-locked Theta and ERN amplitude (Luu et al., 2004; Allen & Trujillo, 2007). On the other hand, the Pe is associated with conscious perception of an error, reflecting motivational aspects and top-down cognitive control (Ridderinkhof et al., 2009). It has been shown that while the ERN is always present following error-trials, the Pe is elicited only in trials in which subjects are aware of their errors (Nieuwenhuis et al., 2001). In the present study, an unexpected correction in the Avatar’s movement was implemented in both the Interactive and Cued conditions. Crucially, only in the Interactive condition one needs to spatially predict the outcome of the partner’s behaviour. Thus, the higher activation of the early error detection system (ERN) in this condition seems to be associated to the need to use information concerning the partner’s movements to guide one’s own movements.

Previous EEG studies have described the occurrence and modulations of ERN and Pe responses to the perception of odd events as well as of errors of a partner in turn taking interactive scenarios (Koban et al., 2010; Kato et al., 2016). Similarly, other studies have investigated the occurrence of ERN and Pe responses during interpersonal musical performance in turn taking set-ups (Maidhof et al., 2010; Huberth et al., 2018). One EEG study demonstrates that performing errors together with a partner modulates neural activity related to outcome evaluation (i.e. Feedback-Related Negativity is larger for joint errors compared to other’s ones) but has less impact on activity related to the motivation to adapt future behaviour (i.e. P3b is not modulated by own, joint or other’s errors; Loehr et al., 2015). This suggests that producing a phasic error by pressing the wrong key synchronously with a partner impacts outcome evaluation response rather than generating neural responses associated to adaptive behaviour. Instead, our study characterizes error-related responses when participants need to adapt on-line to the synchronous behaviour of an interactive partner that violates a motor prediction.

### Error-related responses in time-frequency domain

The time-frequency analysis on FCz reveals a greater Theta and Alpha synchronization for the Interactive-Correction condition compared to all other conditions. Although reported separately in the analysis, Theta and Alpha modulation shows a similar and correlated pattern (see Table 2). In previous error-related literature, Theta and Alpha have both been found over fronto-central electrodes during the processing of errors (Pavone et al., 2016, Pezzetta et al., 2018). However, Trujillo and Allen (2007) have argued that activity in the lower Alpha band is due to leakage of the Theta frequency to the neighbouring bands. In our case, the fact that Theta synchronization results show higher increases in power than Alpha and that their activity is correlated (see Table 2) suggests a spread from the Theta band over the Alpha band.

Interestingly, when the Avatar corrected his action the Theta synchronization was reduced in the Cued compared to the Interactive condition suggesting a dissimilar processing of the correction in Interactive and Cued conditions. Furthermore, in the Interactive condition, Theta and Alpha activity were found even in trials when the virtual partner performed no correction. This error-related activity in the absence of error (i.e. Interactive-NoCorrection) suggests that the monitoring system does not only react to unexpected actions but plays a key-role in continuously monitoring the partner’s and ones’ own behaviour in order to integrate the partner’s behaviour when participants’ actions rely on them.

### Frontal theta ERS: Monitoring System and Error Detection

Results in the Theta-band indicate that goal-related and temporal coding of the observed actions might undergo different processing systems. We suggest that the violation of the predicted goal of the observed actions (Correction factor), and the need to adjust to them (Interactive condition), represent the crucial features upon which the error-related monitoring system is based. A parsimonious interpretation of this pattern of results is that the monitoring system is differentially activated by the behavioural relevance of events in the Interactive and Cued conditions. Accordingly, the frontal Theta activity is less present during the Cued condition. In such condition the subject is not engaged in monitoring the partner’s action goals and likely dedicates less resources to processing the partner’s behaviour. Coherently with this, our DICS analysis of the Theta band revealed a fronto-central source estimate (Cohen, 2011; Kovacevic et al., 2012) where the Anterior Cingulate Cortex (ACC) is believed to be a key-part of the cognitive control network.

Theta dynamics shown in the current study provide new insights on the neural underpinnings of cognitive control and action-related processing during motor interactions. Importantly, such an effect was maximal during sudden changes in the virtual partner’s movement. This shows that higher uncertainty in the Interactive condition generates stronger source-located fronto-central Theta.

### Alpha power during interpersonal motor interactions

An often-described EEG marker of engagement in interactive paradigms is the sensorimotor alpha/mu desynchronization over central sites (Tognoli et al., 2007; Dumas et al., 2010; Naeem et al., 2012; Ménoret et al., 2014; Konvalinka et al., 2014; Novembre et al., 2016). This rolandic alpha/mu band activity has been considered an index of the MNS activity since it is suppressed during both action observation and action execution (Cochin et al., 1999; Muthukumaraswamy et al., 2004; Oberman et al., 2005; Pineda, 2005). However, recent studies provide more cautious interpretation concerning the alpha/mu modulation, revising some of the conclusions made about the implication of the MNS in processing self and others’ actions in healthy participants (Coll et al., 2017) and clinical samples - such as people with autism (Dumas et al., 2014). In our analysis, we do not find any modulation of the sensorimotor alpha/mu rhythms between conditions. However even though our time-frequency analysis suggests a spreading of the Theta activity over the Alpha band, we highlight a frontal increase in Alpha power (see Figure 5) for the Interactive-Correction condition. Frontal increase in the Alpha band has been associated with attentional modulation, with the aim of “gating” the incoming flow of information with top-down processing (Benedek et al., 2011) and error monitoring (van Driel et al., 2012).

### Occipito-temporal theta responses during interactions

The action observation network (AON) has been proposed as a neural substrate for action understanding (see for review Rizzolatti et al., 2014; Avenanti et al., 2013; Urgesi et al., 2014). However, recent findings associated the ability to decode an action with activity in the lateral occipito-temporal cortex (Lingnau & Downing, 2015; Tucciarelli et al., 2015). Interestingly, in addition to fronto-central Theta activity associated with error-detection, source analysis revealed activity in the right occipito-temporal cortex. This region is thought to play a role in the processing of body images as indicated by functional methods (Downing et al., 2001; Thierry et al., 2006), virtual lesions (Urgesi et al., 2004; Urgesi et al., 2007) and studies on brain-damaged patients (Moro et al., 2008). More recently we have shown that an occipito-temporal Theta ERS is found during the passive observation of hands and arms images (Moreau et al., 2017) while Tucciarelli et al. (2015) showed that activity in the Theta band over LOTC areas distinguishes between hand pointing and grasping actions. Coherently with this, an fMRI study showed that during the observation of kinematic errors in a reaching task activity of occipito-temporal areas increases parametrically with the dimension of the observed error (Malfait et al., 2010).

Here we describe a Theta increase when the hand of an interactive partner deviates from its expected trajectory. Therefore, we submit that the Theta source detected over occipito-temporal area during Interactive-Correction trials is associated with action re-coding after a deviation from the predicted goal was perceived in the movement of the avatar. This re-coding appears to be a necessary step to adapt to the avatar’s sudden change in movement. This suggests that the increase of Theta activity over occipito-temporal regions during perception of static hand images (Moreau et al., 2017) extends to the perception of dynamic hand movements. Therefore, Theta may be an intrinsic rhythm of occipito-temporal areas that becomes enhanced when attention is deployed over body-related movements (Engel et al., 2001), for example, when the movement of a partner is different from what was predicted. Furthermore, this region seems to be connected to other brain regions as suggested by our connectivity results (PLV), showing an increase of phase-locking between the occipito-temporal and the fronto-central areas. This result suggests that these two regions belong to a common functional neural network recruited during behavioural adaptation in a social context.

### Conclusions

We describe the EEG correlates of error detection during motor interactions with a virtual partner that performed expected or less expected actions. We found that electrocortical markers of error processing were stronger for unpredicted actions; particularly in the Interactive condition during which goal-related and temporal predictions of the partner’s actions are required. Moreover, the source estimates of the Theta frequency markers show the recruitment of fronto-central and occipito-temporal regions, indicating their potential role in processing and integrating visual and motor information during social interactions.

## Acknowledgments

The authors thank Sarah Boukarras, Daniele Esposito, Michele La Sala and Michela Fracassi for helping with the recordings of the data. We also thank Eilidh McCann for proofreading the article. The study was made possible by BrainTrends who provided technical support. BrainTrends did not have any financial or scientific influence on the present study. No conflicts of interest, financial or otherwise, are declared by the authors.

## Supplementary Material

### Behavioural data

We considered the following as behavioural measures: 1) Grasping Synchrony, i.e. the absolute value of the time delay between subjects’ index–thumb contact-times on their bottle and the avatar’s reaching time; 2) Accuracy, that is the number of movements executed correctly (according to the instructions); 3) Reaction Times (RTs), i.e. time from the go-signal to the release of the start button; 4) Movement Times (MTs), i.e. time interval between participants releasing the start button and their index-thumb touching the bottle.

### Motion Kinematics data

Motion tracking was continuously recorded during the experimental blocks. During off-line analyses, the participants’ start button-hand-release times and index-thumb-bottle contact times were used to subdivide the kinematics recordings with the aim of analysing only the reach-to-grasp phase (from start button hand-release to index-thumb contact-times). To obtain specific information on the reaching component of the movement, we analysed wrist trajectory as indexed by the maximum peak of wrist height on the vertical plane (Maximum Wrist Height). To obtain specific information on the grasping component of the movement, we analysed maximum grip aperture (Maximum Grip Aperture, i.e., the maximum peak of index-thumb 3D Euclidean distance). We excluded from the analyses (behavioural, kinematics and EEG) trials in which participants 1) missed the touch-sensitive sensors and thus no response was recorded, 2) released the start button before the go instruction or 3) did not comply with the complementary/imitative instructions.

Behavioural and kinematic values that fell 2.5 SDs above or below each individual mean for each experimental condition were considered as outliers and excluded from the analyses. We calculated the individual mean value in each condition for each of these behavioural and kinematics measures. The obtained values were entered in different within-subject ANOVAs (see below). We used non-parametric tests concerning the Accuracy measures. Kinematics, Accuracy, MTs and RTs results are presented below.

### Additional Analyses and Results

Behavioural, kinematics and EEG (Synchronicity, Reaction Times, Movement Times, Maximum Wrist Height, Maximum Grip Aperture, Theta, Alpha and Beta over FCz) data were analysed through repeated measures ANOVAs; with Correction (Correction, NoCorrection), Condition (Interactive, Cued), Interaction Type (Complementary, Imitative), Movement Type (Precision, Power) as within subject factors. Accuracy was analysed by means of non-parametric tests.

### Behavioural and Kinematics results

#### Synchrony

Because of violations of Normality Assumptions, data were transformed, using logarithmic (log10) transformation.

The ANOVA on Synchrony showed: a significant Correction x Condition x Interaction Type x Movement Type interaction (F(1, 20) = 63.32, p < 0.001), which explained all other significant Main effects and lower level interactions. Post-hoc tests showed that participants were less synchronous when performing Complementary compared to Imitative movements during power grasping in the Interactive Condition; when the Avatar did not correct its movement trajectory (*p* = 0.02). Moreover, performance decreased during correction compared to non-correction trials in the Interactive condition, when performing Complementary movements through Power grips (*p* < 0.001). Performance was worse in Correction trials, during Interactive interactions, when performing Complementary compared to Imitative precision grips (*p* = 0.004). Furthermore, participants were less synchronous in Correction trials, when performing Interactive interactions involving Complementary movements with precision compared to power grips (*p* = 0.001). Moreover, performance decreased during Correction trials in the Interactive compared to the Cued condition, when performing Imitative movements through Precision grips (*p* < 0.001). Finally, performance was worse in Correction trials, when performing Interactive compared to Cued interactions, during Complementary power grips (*p* < 0.001).

#### Reaction Times (RTs)

The ANOVA on Reaction Times showed: a significant Correction x Interaction Type x Movement Type interaction (F(1, 20) = 41.77, *p* < 0.001), which explained all the other significant Main effects and lower level interactions. Post-hoc tests showed that participants were faster to start moving during Correction trials, when performing Complementary actions through Power grips. and during Correction trials, when performing Imitative actions through Precision grips, compared to all the other conditions (all *ps* < 0.001).

#### Movement Times (MTs)

The ANOVA on Movement Times showed a significant main effect of Condition (F(1, 20) = 16.22, *p* < 0.001), indicating that coordinating in the Interactive condition resulted in slower movement times compared to the Cued condition. The ANOVA also showed a significant Condition x Correction interaction (F(1,20) = 19.75, *p* < 0.001). Post-hoc tests showed movement times were slower in Interactive compared to Cued conditions (all ps < 0.001) and in Interactive condition during Correction compared to NoCorrection trials (*p* = 0.02). Moreover, the ANOVA showed a significant Condition x Interaction Type interaction (F(1,20) = 7.78, *p* = 0.02). Post-hoc tests showed movement times were slower in Interactive compared to Cued conditions (all ps < 0.001) and in Interactive condition during Complementary compared to Imitative movements (*p* =0.03). The ANOVA on Movement Times also showed a significant Interaction Type x Movement Type interactions (F(1,20) = 18.16, *p* < 0.001), explained by the higher order Condition x Interaction Type x Movement Type interaction (F(1,20) = 21.38, *p* < 0.001)). Post-hoc tests showed movement times were slower during Correction trials, when performing Complementary movements by means of Power compared to Precision grips (*p* = 0.034) and during Correction trials, when performing Complementary compared to Imitative movements by means of Power grips (*p* < 0.001). Moreover, post-hoc tests showed slower movements times during Correction trials, when performing Imitative compared to Complementary movements by means of Precision grips (*p* = 0.03).

#### Maximum Wrist Height (MaxH)

The ANOVA on Maximum Wrist Height showed a significant Condition x Interaction Type x Movement Type interaction (F(1,20) = 12.98, *p* = 0.001). Post-hoc tests indicated that when performing power grips during the Interactive condition, maximum wrist height was higher during complementary compared to imitative movements (*p* < 0.001). This result highlights the presence of visuo-motor interference between self-executed actions and those observed in the partner as an index of automatic imitation. These results mirror previous studies (Sacheli et al., 2012; 2013; 2015 a; 2015b; Candidi et al., 2015; Curioni et al., 2017), only in the condition during which predictions about the partner’s movements are needed. Visuo-motor interference effects were present only when performing power grips on the lower part of the bottle as, when performing precision grips on the upper part of the bottle, the maximum wrist height is always reached when touching the bottle – thus impossible to modulate.

#### Maximum Grip Aperture (MaxAp)

The ANOVA on Maximum Grip Aperture showed a significant Correction x Condition x Movement Type interaction (F(1, 20) = 147.81, *p* < 0.001), which explained all the other significant Main effects and lower level interactions. Post-hoc tests showed larger maximum grip aperture during Power compared to Precision Grips (all ps < 0.001) and larger maximum grip aperture during Interactive compared to Cued interactions (all ps < 0.001), but not during NoCorrection trials, in Interactive compared to Cued interactions, by means of Power Grips (*p* = 1).

#### Accuracy

A Friedman ANOVA revealed significant cross-condition differences (Chi Sqr. (N = 22, df = 15) = 46.24, *p* < 0.001). Follow-up Wilcoxon Matched Pairs Tests between Correction and NoCorrection conditions showed that Correction condition was more difficult (i.e. less accurate) than NoCorrection condition when performing Complementary-Precision grips in the Interactive Condition (*p* = 0.002, corrected *p* threshold = 0.05/8 = 0.006).

### EEG Analysis

#### Theta over FCz

The ANOVA on Theta synchronization over FCz showed a significant main effect of Correction (F(1, 20) = 49.609, *p* = 0.001) indicating a greater Theta for Correction trials, and a main effect of Condition (F(1, 20) = 93.846, *p* = 0.001) indicating a greater Theta for the Interactive interaction. The ANOVA also revealed a Correction x Condition x Interaction type interaction (F(1, 20) = 12.658, *p* = 0.00257). Post-hoc tests showed larger Theta activity for Correction trials in the Interactive interaction both for complementary and imitative movements compared to the other conditions (*ps* < 0.001), and greater Theta for Correction trials in the Cued interaction when subjects were performing a complementary movement compared to an imitative one (*p* = 0.001). The ANOVA also revealed a Condition x Interaction type x Movement type interaction (F(1, 20) = 6.4726, *p* = 0.018). Post-hoc tests indicated that all NoCorrection trials showed smaller Theta compared to all Correction trials (*ps* < 0.001).

#### Alpha over FCz

The ANOVA on Alpha synchronization over FCz showed a significant main effect of Correction (F(1, 20) = 16.460, *p* = 0.001) indicating a greater Alpha for Correction trials, a main effect of Condition (F(1, 20) = 36.398, *p* = 0.001) indicating a greater Alpha for the Interactive interaction, and a main effect of Movement type (F(1, 20) = 6.8686, *p* = 0.019) indicating a greater Alpha synchronization for power grasps compared to precision ones. The ANOVA also revealed a Correction x Condition interaction (F(1, 20) = 13.6520, *p* = 0.001). Post hoc tests indicated a larger Alpha for Correction trials in the Interactive condition compared to the other conditions (*ps* < 0.001) and a larger Alpha for NoCorrection trials in the Interactive condition compared to Cued ones in both Correction (*p* = 0.001) and NoCorrection (*p* = 0.001).

#### Beta over FCz

The ANOVA on Beta synchronization over FCz showed a significant main effect of Correction F (1, 20) = 20.263, *p* = 0.001) indicating a greater Beta for Correction trials and a main effect of Condition (F(1, 20) = 48.3221, *p* = 0.001) indicating a greater Beta for the Interactive interaction. The ANOVA also revealed a Correction x Condition interaction (F(1, 20) = 22.266, *p* = 0.001). Post hoc tests indicated a larger Beta for Correction trials in the Interactive condition compared to the other conditions (*ps* < 0.00001), and a larger Beta for NoCorrection trials in the Interactive condition compared to Cued ones in both Correction (*p* = 0.001) and NoCorrection (*p* = 0.00001).

A greater Beta synchronization for Correction during the Interactive condition might be linked to the so-called Beta rebound, associated with the degree of error in a movement (Tan et al., 2014).

